# Environmental drivers of body size in North American bats

**DOI:** 10.1101/2021.07.28.454183

**Authors:** J.M. Alston, D.A. Keinath, C.K.R. Willis, C.L. Lausen, J.M. O’Keefe, J.D. Tyburec, H.G. Broders, P.R. Moosman, T.C. Carter, C.L. Chambers, E.H. Gillam, K. Geluso, T.J Weller, D.W. Burles, Q.E. Fletcher, K.J.O. Norquay, J.R. Goheen

## Abstract

Bergmann’s Rule—which posits that larger animals live in colder areas—is thought to influence variation in body size within species across space and time, but evidence for this claim is mixed. We tested four competing hypotheses for spatio-temporal variation in body size within bat species during the past two decades across North America. Bayesian hierarchical models revealed that spatial variation in body mass was most strongly (and negatively) correlated with mean annual temperature, supporting the heat conservation hypothesis (historically believed to underlie Bergmann’s Rule). Across time, variation in body mass was most strongly (and positively) correlated with net primary productivity, supporting the resource availability hypothesis. Climate change could influence body size in animals through both changes in mean annual temperature and in resource availability. Rapid reductions in body size associated with increasing temperatures have occurred in short-lived, fecund species, but such reductions likely transpire more slowly in longer-lived species.

## 1 Introduction

Body size influences every aspect of organismal biology, including lifespan (Lindstedt & Calder 1981; Speakman 2005), metabolism (Brown *et al*. 2004; Clarke *et al*. 2010), movement rates (Jetz *et al*. 2004; Noonan *et al*. 2020), reproductive biology (Fenchel 1974; Blueweiss *et al*. 1978), and extinction risk (Brown 1995; Ripple *et al*. 2017). Understanding the factors that drive variation in body size is thus among the most important goals in ecology (Kaspari 2005). Bergmann’s Rule (Bergmann 1847; Salewski & Watt 2017), which states that animals residing in colder climates are larger than those residing in warmer climates, is a widely known macroecological pattern. Although originally and primarily applied to differences in body size between closely related species, Bergmann’s Rule is often believed to extend to differences in body size within species as well (Ashton 2002; Meiri & Dayan 2003; Blackburn & Hawkins 2004; Watt *et al*. 2010; Riemer *et al*. 2018).

The mechanism traditionally hypothesized to underlie Bergmann’s Rule is that increased size facilitates body heat conservation (hereafter, the “heat conservation hypothesis”; Bergmann 1847; Mayr 1956; Ashton 2002; Watt *et al*. 2010). To maintain a stable, elevated body temperature, homeotherms must operate at higher metabolic rates when faced with cooler ambient temperatures, thus losing substantial metabolic heat to the environment (McCafferty *et al*. 2011; Fristoe *et al*. 2015). The ratio between surface area and volume decreases with increasing body size, so while absolute heat loss increases with increasing body size, smaller animals require greater metabolic heat production to offset heat loss across their relatively large surface areas (Withers *et al*. 2016). Larger body size could therefore be an adaptation to climates with cooler average temperatures.

Despite its intuitive appeal, empirical support for the heat conservation hypothesis within species is mixed. Although ecologists have accumulated substantial evidence that individuals within species tend to be larger in colder climates (e.g., Smith *et al*. 1995; Ashton 2002; Meiri & Dayan 2003), more recent and more comprehensive tests have failed to find consistent relationships between temperature and the body sizes of individuals within species (Riemer *et al*. 2018). Additionally, physiologists have questioned the validity of the heat conservation hypothesis on physiological grounds (Scholander 1955; McNab 1971; Geist 1987). In sum, and despite the widespread acceptance of the heat conservation hypothesis in the ecological literature, the extent to which variation in average temperature translates to variation in body size within species remains an open question.

Skepticism surrounding the primary mechanism assumed to underlie Bergmann’s Rule has resulted in other hypotheses for geographical clines in body size within species that are consistent with Bergmann’s Rule (Meiri *et al*. 2007; Salewski & Watt 2017). For example, larger individuals exhibit acute heat stress at lower temperatures than smaller individuals, and thus experience greater risk of mortality from heat stress than smaller individuals (hereafter the ‘heat mortality hypothesis’; Smith *et al*. 1995; Peralta-Maraver & Rezende 2021; but see McKechnie *et al*. 2021). This idea posits an additional (or alternative) mechanism by which animals from warmer climates are smaller than their counterparts from colder climates, and is supported in the genus *Neotoma* (i.e., woodrats; Brown & Lee 1969; Smith *et al*. 1995). A second alternative is the ‘resource availability hypothesis’, through which increased resource availability—often correlated with temperature across the globe (Gillman *et al*. 2015; Chu *et al*. 2016)—results in larger individuals and drives biogeographical patterns in body size (e.g., Rosenzweig 1968; McNab 2010; Huston & Wolverton 2011; Yom-Tov & Geffen 2011; Kelly *et al*. 2018). If true, clinal variation in body size consistent with Bergmann’s Rule could arise over limited geographic extents (e.g., an elevational gradient where increased precipitation increases productivity as temperature decreases), but body sizes should *decrease* as temperatures decrease at very large (i.e., continental) spatial scales (which would contradict Bergmann’s Rule). Finally, a third hypothesis proposed to explain Bergmann’s Rule is the ‘starvation resistance’ (or ‘seasonality’) hypothesis. According to this hypothesis, large body size buffers against resource scarcity driven by seasonality (Boyce 1979). Because seasonality increases at higher latitudes and fasting endurance decreases at colder temperatures (Lindstedt & Boyce 1985), this dynamic may produce a size cline consistent with Bergmann’s Rule. The starvation resistance hypothesis has received support from studies on songbirds (Jones *et al*. 2005), muskrats (*Ondatra zibethicus*; Boyce 1978), and bobcats (*Lynx rufus*; Wigginton & Dobson 1999).

Some ecologists and evolutionary biologists have suggested that Bergmann’s Rule should apply over time as well as space (e.g., Smith *et al*. 1995; Van Buskirk *et al*. 2010; Merckx *et al*. 2018; Weeks *et al*. 2020). In other words, as temperatures fluctuate over time, the average size of individuals within a species should decrease as temperatures rise and should increase as temperatures fall. Although early studies of this temporal equivalent to Bergmann’s Rule focused on time scales of thousands of years (Smith *et al*. 1995), more recent studies have found that changes in body size can occur over decades or even years (Van Buskirk *et al*. 2010; Weeks *et al*. 2020). However, and similar to the original (spatial) conceptualization of Bergmann’s Rule, empirical evidence for this temporal equivalent is mixed (Sheridan & Bickford 2011; Yom-Tov & Geffen 2011; Teplitsky & Millien 2014), perhaps because other mechanisms—analogous to alternative mechanisms for Bergmann’s Rule detailed above—influence shifts in body size over time. For example, extreme climatic events can trigger rapid evolution of traits (Campbell-Staton *et al*. 2017; Donihue *et al*. 2018), which is consistent with the heat mortality and starvation resistance hypotheses. As another example, positive effects of periods of high resource availability on fat reserves and growth have been documented in many taxa (e.g., Brett 1971; Boutin & Larsen 1993; Altmann & Alberts 2005; Monteith *et al*. 2014), and would support the resource availability hypothesis. Testing these alternative hypotheses across both space and time provides a lens through which to anticipate how changes in climate may affect body size in the future, as well as the pace at which changes in body size may occur.

To evaluate the mechanistic underpinnings of Bergman’s Rule, we tested whether spatial and temporal variation in body mass of North American bats is best supported by the heat conservation, heat mortality, resource availability, or starvation resistance hypotheses (summarized in Table 1). As with many taxa, the intraspecific formulation of Bergmann’s Rule is exhibited by some species of bats (e.g., Burnett 1983; Bogdanowicz 1990; Lausen *et al*. 2008, 2019), but does not appear to be the norm among the clade as a whole (Riemer *et al*. 2018). Critically, extensive records of bat captures permit a rare opportunity to test for Bergmann’s Rule and evaluate its associated hypotheses while accounting for other factors (e.g., sex, age, reproductive condition, and time of year) that influence body size. We compiled 17 such data sets and used Bayesian hierarchical models to weigh evidence for each hypothesis across both space and time for 20 species of North American bats. We expected observed patterns of variation in body mass to be driven by the same process or processes across both time and space. In other words, if variation in body mass across space was best explained by one of our four hypotheses, we also expected variation in body mass across time to be best explained by the same hypothesis. This would provide strong evidence for a consistent selective force driving variation in body size.

**Table 1.** Descriptions of the hypotheses tested in this paper, including the name, spatial version of the hypothesis, proxy data used to test the spatial hypothesis, temporal version of the hypothesis, and proxy data used to test the temporal hypothesis.

| **Hypothesis Name** | **Spatial Hypothesis** | **Spatial Proxy Data** | **Temporal Hypothesis** | **Temporal Proxy Data** |
| --- | --- | --- | --- | --- |
| *Heat Conservation* | Because larger body size increases an individual's ability to conserve body heat, individuals will be larger in areas where average temperatures are lower. | Mean annual temperature (1970-2000; WorldClim; Fick and Hijmans 2017) | Because larger body size increases an individual's ability to conserve body heat, individuals will be larger after years in which average temperatures are lower. | Mean temperature from April 1 of capture year until date of capture (DAYMET; Thornton et al. 2020) |
| *Heat Mortality* | Because larger individuals tend to have lower critical thermal maxima, individuals will be smaller in areas where maximum temperatures are higher. | Mean annual maximum temperature (1980-2010; DAYMET; Thornton et al. 2020) | Because larger individuals tend to have lower critical thermal maxima, individuals will be smaller after years in which maximum temperatures are higher. | Maximum temperature in the preceding 365 days (DAYMET; Thornton et al. 2020) |
| *Resource Availability* | Because individuals living in more productive environments tend to be larger, individuals will be larger in areas where primary productivity is higher. | Net primary productivity during April – October (2000-2016; MODIS; Stockli 2020) | Because individuals living in more productive environments tend to be larger, individuals will be larger after years in which primary productivity is higher. | Net primary productivity in months preceding capture, inclusive of month of capture (MODIS; Stockli 2020) |
| *Starvation Resistance* | Because larger body size increases an individual's ability to survive periods of resource scarcity, individuals will be larger in areas where periods of resource scarcity (e.g., winters) are most severe. | Average minimum temperature in April, May, September, and October (1970-2000; WorldClim; Fick and Hijmans 2017) | Because larger body size increases an individual's ability to survive periods of resource scarcity, individuals will be larger after years in which periods of resource scarcity (e.g., winters) are most severe. | Average minimum temperature in the April, May, September, and October preceding capture (DAYMET; Thornton et al. 2020) |

## 2 Methods

### 2.1 Data Collection

We compiled biometric data on bats captured throughout North America using mist nets between 2000 and 2016 (Fig. 1). All biometric data contained information on capture location, date of capture, species, sex, age class, reproductive state, and mass. Because body mass varies with species, sex, age class, reproductive state, and time of year, we accounted for potential variation related to these factors by calculating the mean mass for each species/sex/reproductive state combination in each month, subtracting the corresponding mean value from the mass of each individual in the data set, and dividing this by the standard deviation of body mass values for that species. The final data set included only data from adult bats captured between April and October, for species represented by ≥ 150 individuals and that were captured across ≥ 2.5° of latitude.

**Fig. 1.**
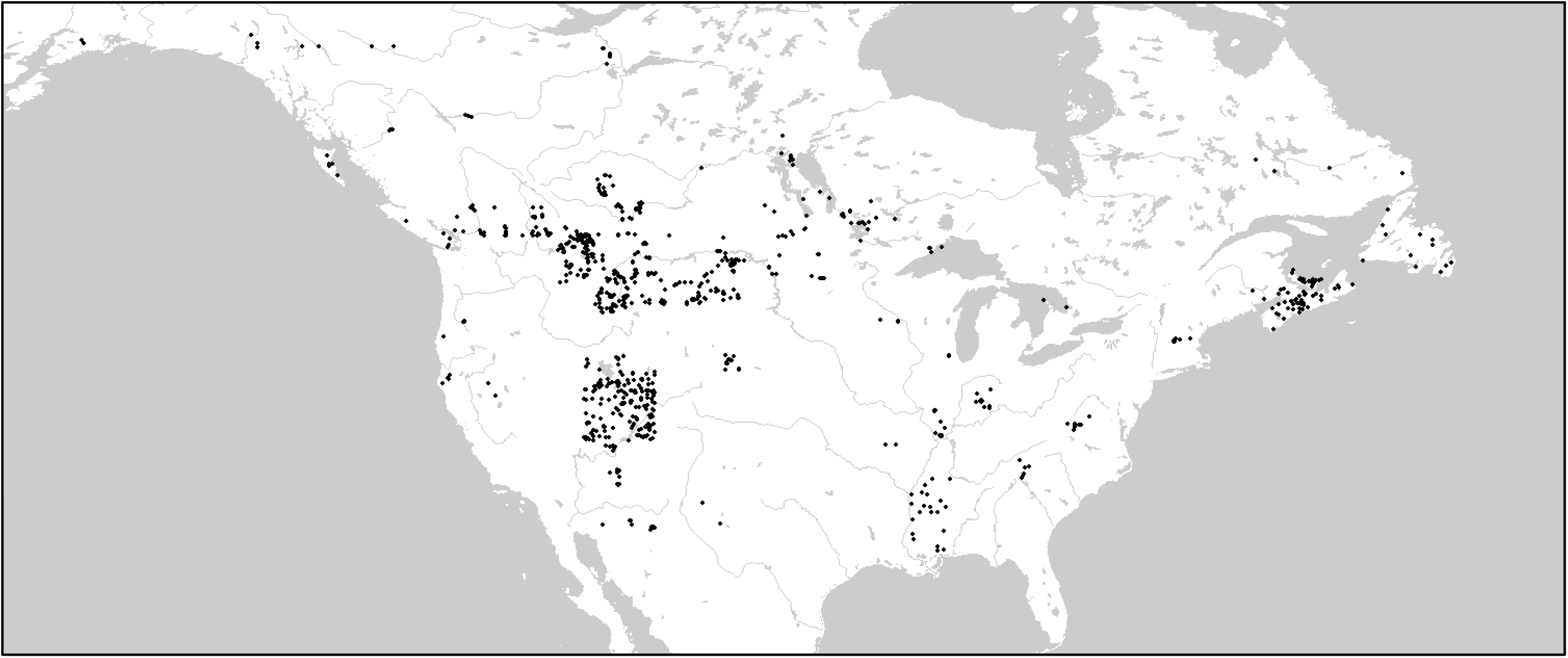
Map of capture locations for bats included in our analyses. Our final data set included 31,303 bats sampled from 1,190 sites along a >30° gradient in latitude.

To test hypotheses for clinal variation in body mass, we extracted environmental variables from remotely sensed raster data sets. To test the heat conservation hypothesis across space, we extracted data for each capture location from the 30-second (∼1 km) resolution version of the WorldClim 2.1 mean temperature data set (mean annual temperature, 1970-2000; Fick & Hijmans 2017). We centered mean annual temperatures in our data set at zero by subtracting the mean annual temperature across all capture locations. To test the heat conservation hypothesis across time, we extracted data for each capture location from the DAYMET daily climate summaries 1- km resolution data set (Thornton *et al*. 2020) using the ‘daymetr’ package (version 1.4; Hufkens *et al*. 2018). We used those data to calculate the midpoint of low and high temperatures across all days from 1 April until the date each bat was captured, and then subtracted the average of this value at the capture location during our study period (2000-2016) to obtain a final centered metric of year-to-year differences in mean temperatures. This represents roughly the period in which a bat would be active in a given year (dates before 1 April are likely to be spent in hibernation or in winter ranges).

To test the heat mortality hypothesis across space, we extracted data for each capture location from the DAYMET daily climate summaries 1-km resolution data set (Thornton et al. 2020) and used those data to calculate the maximum temperature at each capture location in each year between 1980 and 2010 (the earliest 30-year period available). We then calculated the mean annual maximum temperature across this 30-year period at each site and subtracted the mean annual maximum temperature across all sites to obtain a final centered metric of long-term maximum annual temperatures. To test the heat mortality hypothesis across time, we extracted data for each capture location from the DAYMET daily climate summaries 1-km resolution data set (Thornton et al. 2020). We calculated the maximum temperature in the prior 365 days for each capture event, then subtracted the long-term average for this value at the site of capture to calculate a final centered metric of year-to-year differences in maximum temperatures.

To test the resource availability hypothesis across space, we extracted data for each capture location from the 0.1-degree (∼10 km) resolution version of the MODIS monthly net primary productivity data set (Stockli 2020). Primary productivity is positively correlated with insect biomass across both space (Borer *et al*. 2012; Lind *et al*. 2017) and time (Bell 1985; Frith & Frith 1985), and summer precipitation—another common proxy for resource availability—is positively correlated with annual survival in little brown bats (*Myotis lucifugus*; Frick *et al*. 2010). We averaged monthly net primary productivity across months during the active season for bats (April-October) for all available years (2000-2016), then divided by the mean value across all sites to obtain a final metric centered at one. To test the resource availability hypothesis across time, we extracted data from the same rasters and averaged net primary productivity for months preceding the date a bat was captured (in the year of capture, inclusive of the month of capture, starting in April), then divided by the average of this value at the site of capture for the entire time period.

To test the starvation resistance hypothesis across space, we extracted data for each capture location from the 30-second (∼1 km) resolution version of the WorldClim 2.1 minimum temperature data set (mean minimum temperature, 1970-2000; Fick and Hijmans 2017). To estimate the severity of resource limitation in the period in which bats are most resource-limited, we averaged minimum temperatures across the months of September, October, April, and May, which roughly represent night-time temperatures during the times of year when bats tend to be most energetically vulnerable. Regardless of whether they hibernate or migrate for the winter, bats at temperate latitudes must gain a substantial amount of weight in the autumn (Kunz *et al*. 1998; Lacki *et al*. 2015; Guglielmo 2018; Cheng *et al*. 2019; Sommers *et al*. 2019), and they tend to be energetically stressed in the early spring before insects become abundant (Arlettaz *et al*. 2001; Encarnação *et al*. 2004; Jonasson & Guglielmo 2019). Daily minimum temperatures during autumn and spring thus represent a biologically informed proxy for resource limitation. We centered mean minimum spring and autumn temperatures in our data set at zero by subtracting the mean minimum spring and autumn temperature across all capture locations. To test the starvation resistance hypothesis across time, we extracted data from the DAYMET daily climate summaries 1-km resolution data set (Thornton et al. 2020). We averaged the minimum daily temperatures for the spring (April and May) and autumn (September and October) preceding the date on which a bat was caught, and subtracted the average value at the site of capture during our study period.

### 2.2 Statistical Analysis

We used the R statistical software environment (version 4.0.2; R Core Team 2020) to quantify the influence of our environmental variables on bat body mass across both space and time. We used the modelling software ‘Stan’ (Carpenter *et al*. 2017) via the R package ‘brms’ (version 2.13.3; Bürkner 2017) to build a single Gaussian-family Bayesian model for each species (i.e., 20 models in total; Fig. A1) to quantify the effects on body mass of the environmental predictors detailed above. Each model included 3 chains that were run for 12,000 iterations (2,000 iterations of warm-up and 10,000 iterations of sampling). We assessed chain convergence using the Gelman-Rubin diagnostic (Ȓ) and precision of parameter estimation using effective sample size. Ȓ < 1.01 and effective sample sizes > 10,000 represent acceptable convergence and parameter precision (Gelman *et al*. 2013; Kruschke 2015). We used leave-one-out cross-validation to check model fit using the R packages ‘loo’ (version 2.3.1; Vehtari *et al*. 2017), and ‘bayesplot’ (version 1.7.2; Gabry *et al*. 2019) to visually assess the cross-validated probability integral transform.

## 3 Results

The final data set contained 31,303 individuals of 20 species captured at 1,190 locations (Fig. 1; Table A1). Most species were larger at higher latitudes, but body size remained relatively constant over our study period (Fig. A2). Significant spatial and temporal variation existed among all predictor variables, enabling detection of meaningful relationships between body mass and predictor variables (Fig. A3).

### Spatial Variation in Body Mass

Spatial variation in body mass most strongly supported the heat conservation hypothesis, with most species exhibiting greater body mass in areas with colder mean annual temperatures (Fig. 2A). For 15 of 20 species, body mass declined with increasing mean annual temperature (i.e., β < 0), and the probability that the coefficient was less than zero was >95% for 6 of these species (*Myotis lucifugus*, *Eptesicus fuscus*, *Lasionycteris noctivagans*, *M. ciliolabrum*, *M. evotis*, and *Parastrellus hesperus*). Most species exhibited minimal variation in body mass with respect to maximum temperature (Fig. 2B), primary productivity (Fig. 2C), and spring/autumn temperatures (Fig. 2D), suggesting a lack of support for the heat mortality, resource availability, and starvation resistance hypotheses, respectively. For these three hypotheses, coefficients were relatively evenly distributed around 0; 90% credible intervals overlapped 0 in most cases, and credible intervals that did not overlap zero were distributed relatively evenly around zero.

**Fig. 2.**
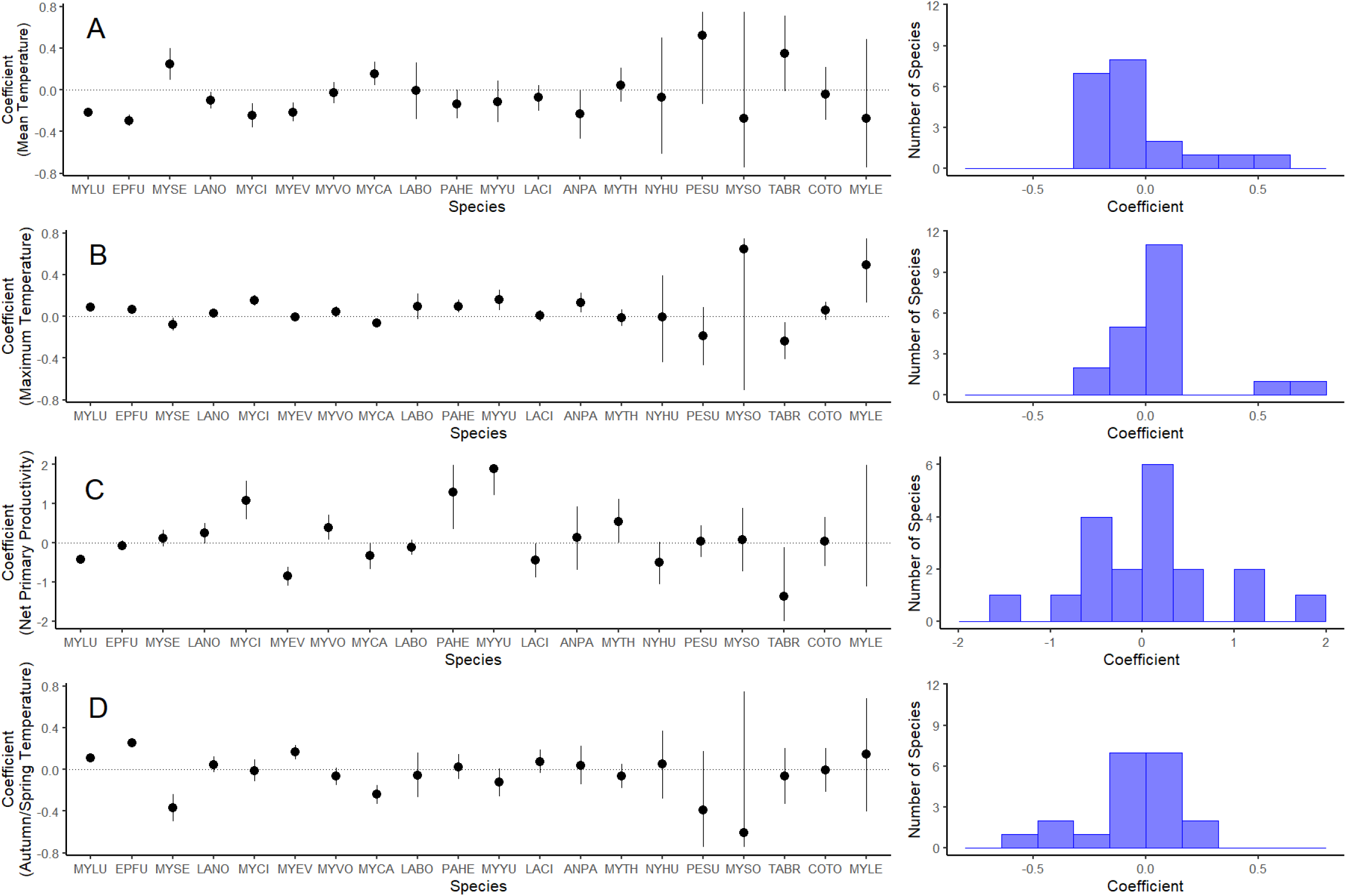
Intraspecific patterns in body mass across space in 20 species of North American bats, which most strongly support the heat conservation hypothesis. In the left column, we plotted the regression coefficient (slope) for each species’ relationship between body mass and the predictor variable of interest (points) and 90% credible intervals (lines). Points above the dotted line at 0 indicate species in which individual body mass increased as the variable of interest (A. Mean annual temperature; B. Maximum annual temperature; C. Autumn/spring temperature; D. Net primary productivity) increased. Species are ordered from largest (left) to smallest (right) sample sizes. In the right column, we plotted histograms of the coefficients. Row A represents tests of the heat conservation hypothesis, Row B represents tests of the heat mortality hypothesis, Row C represents tests of the resource availability hypothesis, and Row D represents tests of the starvation resistance hypothesis. Distributions centered on zero indicate no consistent effect of the variable of interest on body mass, while distributions centered asymmetrically around zero indicate directional effects. Credible intervals were truncated at the limit of the y-axis for ease of interpretation. The mean estimate of the coefficient for the effect of net primary productivity on body mass for *Myotis leibii* (MYLE; 4.29) was excluded from the y-axis of that graph to improve interpretability of coefficient estimates for the other species, but the 90% credible interval for that estimate crosses zero as shown in the graph. Species codes are listed in Table A1.

### Temporal (Interannual) Variation in Body Mass

Temporal variation in body mass most strongly supported the resource availability hypothesis, with most species exhibiting greater body mass during years in which net primary productivity was higher (Fig. 3C). For 14 of 20 species, body mass increased with increasing net primary productivity (i.e., β > 0), and the probability that the coefficient was above zero was >95% for 7 of these species (*Myotis lucifugus*, *Eptesicus fuscus*, *M. ciliolabrum*, *M. californicus*, *Perimyotis subflavans*, *M. sodalis*, *M. leibii*). Most species exhibited little variation in body mass with respect to year-to-year differences in mean annual temperatures (Fig. 3A), maximum temperatures (Fig. 3B), or spring/autumn temperatures (Fig. 3D), suggesting a lack of support for the heat conservation, heat mortality, and starvation resistance hypotheses, respectively. For these hypotheses, coefficients were relatively evenly distributed around 0, 90% credible intervals overlapped 0 in most cases, and credible intervals that did not overlap zero were relatively evenly distributed around zero or were distributed in the direction opposite most coefficients.

**Fig. 3.**
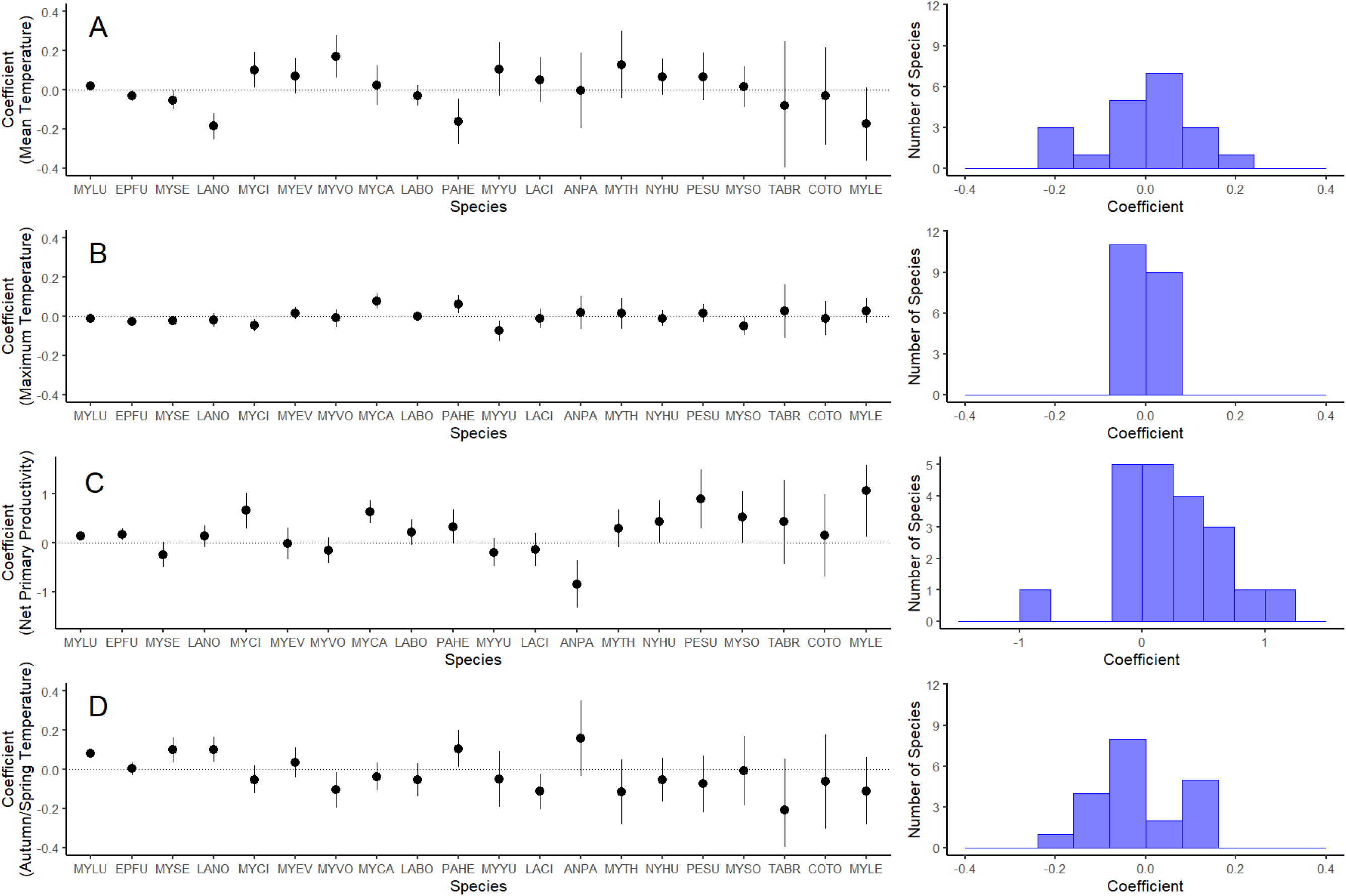
Intraspecific patterns in body mass across time in 20 species of North American bats, which most strongly support the resource availability hypothesis. In the left column, we plotted the regression coefficient (slope) for each species’ relationship between body mass and the predictor variable of interest (points) and 90% credible intervals (lines). Points above the dotted line at 0 indicate species with larger masses as the variable of interest (A. Mean annual temperature; B. Maximum annual temperature; C. Autumn/spring temperature; D. Net primary productivity) increased. Species are ordered from largest (left) to smallest (right) sample sizes. In the right column, we plotted histograms of the coefficients. Row A represents tests of the heat conservation hypothesis, Row B represents tests of the heat mortality hypothesis, Row C represents tests of the resource availability hypothesis, and Row D represents tests of the starvation resistance hypothesis. Distributions centered on zero indicate no consistent effect of the variable of interest on body mass, while distributions centered asymmetrically around zero indicate consistent effects. Credible intervals were truncated at the limit of the y-axis for ease of interpretation. Species codes are listed in Table A1.

## 4 Discussion

Although intraspecific clines in body size have received attention for nearly two centuries (Bergmann 1847; Watt *et al*. 2010), recent studies have cast doubt on both their prevalence and the generality of the mechanisms underlying them (e.g., Meiri *et al*. 2007; Muñoz *et al*. 2014; Freeman 2017; Riemer *et al*. 2018). We used North American bats as a model system to test four competing hypotheses (Table 1) for intraspecific variation in body mass consistent with Bergmann’s Rule, a well-known macroecological pattern. Further, because recent evidence suggests that the mechanisms underlying such geographical clines could be causing rapid evolutionary change in body size (e.g., Van Buskirk et al. 2010; Merckx et al. 2018; Weeks et al. 2020), we also tested the ability of these four hypotheses to describe interannual variation in body mass. Although no hypothesis described variation in body mass across every species, spatial variation in body mass of bats was most consistently correlated with mean annual temperature (supporting the heat conservation hypothesis, originally assumed to underlie Bergmann’s Rule), and temporal variation in body mass was most consistently correlated with net primary productivity (supporting the resource availability hypothesis). In tandem, our results highlight that both spatial and temporal patterns of variation in body size have an energetic basis, but via two distinct pathways: spatial variation in body size is driven by energy loss to the environment in the form of heat, and temporal variation in body size is driven by energy gain from abundant food.

Across North America, body mass of bats was most strongly correlated with mean annual temperature, matching the traditional hypothesis—the heat conservation hypothesis—for Bergmann’s Rule (Bergmann 1847; Mayr 1956). However, this mechanism had little influence on temporal variation in body size, perhaps because selective pressure via size-dependent differences in energy expenditure could take considerable time to manifest. Compared to the heat mortality and starvation resistance hypotheses—which assume punctuated bouts of high mortality driven by extreme heat and severe resource scarcity, respectively—the heat conservation hypothesis posits more gradual selection on body size. Differences in survival and reproduction between small individuals and large individuals may therefore fail to manifest in measurable population-level variation in body size, even after unusually warm or cold years. Only after climate departs from historical norms over many generations should body size change at the population level.

Recent research has cast doubt on the assumption that the heat conservation hypothesis underlies Bergmann’s Rule (Riemer et al. 2018). Using museum specimens of a wide array of endotherms (including bats) collected across the globe, Riemer et al. (2018) found that intraspecific variation in body mass did not vary with mean annual temperature. Our results contradict this finding, likely because we were able to account for confounding sources of variation in body mass (e.g., sex, reproductive status, time of year, resource availability). A diverse array of factors contributes to variation in body mass, and their cumulative influence could swamp variation driven by mean annual temperature (Jones *et al*. 2005; Meiri *et al*. 2007; Nunes *et al*. 2017). Given this challenge, carefully accounting for potential confounding factors is necessary for clarifying the extent to which mean annual temperature drives variation in body size within species. Additionally, our threshold for minimum sample sizes was higher than the threshold used by Riemer et al. (*n* = 150 vs. *n* = 30), and our ability to detect strong evidence for our best-supported hypotheses was positively correlated with sample size. Of the coefficients with > 95% probability of supporting the best-supported hypotheses, 5 of 6 coefficients that supported the heat conservation hypothesis across space and 4 of 7 coefficients that supported the resource availability process across time were located in species with *n* > 1,000 individuals. Given the degree of variation inherent in such broadly collected data, compiling very large data sets (*n* > 1,000) may be necessary to detect Bergmann’s Rule within species in wild populations.

That resource availability might drive body mass variation temporally but not spatially is consistent with predictions of the ideal free distribution model of resource selection (Fretwell & Lucas 1969; Royama 1970) and the “more-individual hypothesis” for species-energy relationships (Wright 1983; Srivastava & Lawton 1998; Storch *et al*. 2018). If individuals within a species are distributed in an ideal free manner, populations should be denser in areas with greater resource availability, such that *per capita* resource availability is roughly equivalent over their geographic range. In this scenario, individuals should not necessarily be appreciably larger or heavier in resource-rich areas than in resource-poor areas, but populations should be denser or sparser, respectively. In other words, additional energy is converted into additional individuals, rather than larger individuals. However, if resource availability changes from year to year, this equilibrium can be disrupted, leading to temporary situations in which *per capita* resource availability is higher in some areas than others until population densities reach a steady state of resource availability. In this scenario, individuals would likely be larger or heavier in (temporarily) resource-rich areas than in (temporarily) resource-poor areas, and this temporal variation in body mass would be driven more by changes in nutritional condition (i.e., fat reserves and muscle mass) than by differences in body size arising from directional selection. This dynamic is likely to be particularly pronounced in long-lived species that produce few offspring (such as bats; Wilkinson & South 2002), because population density cannot rapidly track changes in resource availability via increases in recruitment.

Importantly, our analyses indicate that the processes that drive spatial patterns in body size might not produce equivalent temporal patterns over ecological time scales. Variation in body size occurs both temporally and spatially, but the underlying processes are likely distinct and could manifest over markedly different timescales. Motivated by patterns of spatial variation in body size, many biologists have attempted to quantify analogous patterns through time, typically over the course of years or decades (e.g., Sheridan & Bickford 2011; Caruso *et al*. 2014; Teplitsky & Millien 2014). However, spatial patterns could take centuries or millennia to arise, even when they are relatively clear-cut (and spatial patterns in body size are rarely so). This is especially true for long-lived species, for which the pace of change is likely to be slower than for short-lived, more fecund species.

Climate change will likely induce changes in body size for animals, but such changes may be more complex than has been appreciated. Over the nearly 2 decades that we collected data, the primary driver of short-term (annual) variation in body size was resource availability. Increases in mean annual temperatures could make many ecosystems more productive for a longer portion of the year, but changes in precipitation can both accentuate and dampen such shifts in productivity (Chu *et al*. 2016; La Pierre *et al*. 2016). Any changes in body size driven by climate change will therefore depend on the extent to which mean annual temperature, amount of precipitation, and timing of precipitation are altered for a given area. Moreover, and because net primary productivity does not meaningfully influence body size across space, any such changes are likely to be transient, renormalizing over time if humans eventually curb greenhouse gas emissions.

Life history traits should mediate the influence of climate change on body size. The most compelling evidence of rapid changes in body size due to climate change comes from songbirds (e.g., Van Buskirk *et al*. 2010; Weeks *et al*. 2020), which are shorter lived and more fecund than bats. Controlling for size, bats live on average 3.5 times longer than other placental mammals (Wilkinson & South 2002). Individuals with lifespans > 30 years have been documented in several species, and bats typically produce only 1-2 offspring per year (Wilkinson & South 2002; Barclay & Harder 2003). Because pace of life is positively correlated with the pace of evolution (i.e., smaller, more fecund species tend to evolve more rapidly; Martin & Palumbi 1993; Gillooly *et al*. 2005; Nabholz *et al*. 2008), the processes that lead to spatial variation in body size should arise faster over time in short-lived species than in long-lived species. Additional studies that enable direct comparisons of the pace of body size change across taxa with different life history strategies will increase ecologists’ understanding of the extent and pace of changes in body size caused by climate change.

While recent evidence indicates that higher temperatures caused by climate change are inducing rapid evolution in body size in some species (Van Buskirk *et al*. 2010; Gardner *et al*. 2011; Weeks *et al*. 2020), we found no evidence that this is occurring in bats. Spatial variation in body mass of North American bats is consistent with the heat conservation hypothesis for Bergmann’s Rule, but the heat conservation hypothesis does not explain variation in body size over time. Instead, temporal variation in body mass over the past two decades appears to be driven largely by year-to-year changes in resource availability. For bats and other long-lived species, temperature-induced reductions in body size could take substantially longer to manifest than for short-lived, more fecund species, and will be obscured by variation in resource availability.

## Acknowledgements

Many thanks to J.A. Rick for help with analytical code and helpful comments on early versions of this manuscript. We thank D. Bachen, D. Blouin, S. Bradley, M.-A. Collis, K. Cross, N. Dorville, H. Gates, K.N. Geluso, J. Huebschman, K. Jonasson, D. Nagorson, T. Snow, D. Sparks, H. Thomas, J. Veilleux, B. Walters, J. Whitaker, many Utah Division of Wildlife biologists and technicians, and many others for their work to collect and compile data used for this project. We also thank the Alberta Conservation Association, Arizona Biomedical Research Commission, Arizona Game and Fish Department Heritage Fund, Arizona Game and Fish Department State Wildlife Grant, Bat Conservation International, British Columbia Hydro and Power Authority, British Columbia Ministry of Environment, British Columbia Fish and Wildlife Compensation Program, Habitat Conservation Trust Foundation, Columbia Basin Trust, Indiana Department of Natural Resources, Montana Natural Heritage Program Core Fund, National Science Foundation, Natural Sciences and Engineering Research Council of Canada, Nebraska Game and Parks Commission, North Dakota Department of Agriculture, North Dakota Game and Fish Department, Northern Arizona University, Parks Canada, State of Arizona Technology and Research Initiative Fund, University of Wyoming College of Arts and Sciences, University of Wyoming Department of Zoology and Physiology, US Bureau of Land Management, US Department of Agriculture Wildlife Services, US Department of Agriculture Natural Resources Conservation Service, US Department of Defense Legacy Resource Management Program, US Fish and Wildlife Service, US Forest Service, US Forest Service Rocky Mountain Research Station, US Forest Service Southern Research Station, US Geological Survey, US National Park Service, Utah Division of Wildlife, Utah Endangered Species Mitigation Fund, Virginia Department of Wildlife Resources, and funders of Wildlife Conservation Society Canada’s Western Canada Bat Conservation Program (wcsbats.ca) for funding that made this project possible. This paper includes data from the Hardwood Ecosystem Experiment, a partnership of the Indiana Department of Natural Resources, Purdue University, Ball State University, Indiana State University, Drake University, and The Nature Conservancy. This work was partly funded by the Center for Advanced Systems Understanding (CASUS) which is financed by Germany’s Federal Ministry of Education and Research (BMBF) and by the Saxon Ministry for Science, Culture and Tourism (SMWK) with tax funds on the basis of the budget approved by the Saxon State Parliament.

## Appendix 1 Supplementary Materials

**Table A1.** Table including species included in our analysis, species code (used in figures), the number of individuals included in each species’ model, the latitudinal range covered by individuals of each species (in degrees), and the number of distinct site-year combinations at which each species was captured.

| Species | Species Code | Number of Individuals | Latitudinal Range | Number of Site-years |
| --- | --- | --- | --- | --- |
| *Antrozous pallidus* | ANPA | 567 | 17.50 | 136 |
| *Corynorhinus townsendii* | COTO | 234 | 18.32 | 92 |
| *Eptesicus fuscus* | EPFU | 4,123 | 28.29 | 551 |
| *Lasiurus borealis* | LABO | 918 | 18.55 | 132 |
| *Lasiurus cinereus* | LACI | 824 | 20.82 | 236 |
| *Lasionycteris noctivagans* | LANO | 1,572 | 20.73 | 300 |
| *Myotis californicus* | MYCA | 1,183 | 21.98 | 212 |
| *Myotis ciliolabrum* | MYCI | 1,540 | 20.07 | 318 |
| *Myotis evotis* | MYEV | 1,353 | 21.17 | 384 |
| *Myotis leibii* | MYLE | 150 | 8.11 | 67 |
| *Myotis lucifugus* | MYLU | 12,293 | 23.53 | 693 |
| *Myotis septentrionalis* | MYSE | 1,702 | 25.39 | 243 |
| *Myotis sodalis* | MYSO | 285 | 2.54 | 48 |
| *Myotis thysanodes* | MYTH | 495 | 18.90 | 159 |
| *Myotis volans* | MYVO | 1,309 | 24.38 | 307 |
| *Myotis yumanensis* | MYYU | 847 | 18.24 | 136 |
| *Nycticeius humeralis* | NYHU | 403 | 13.59 | 59 |
| *Parastrellus hesperus* | PAHE | 905 | 8.73 | 129 |
| *Perimyotis subflavans* | PESU | 339 | 14.50 | 86 |
| *Tadarida brasiliensis* | TABR | 261 | 10.50 | 59 |

**Fig. A1.**
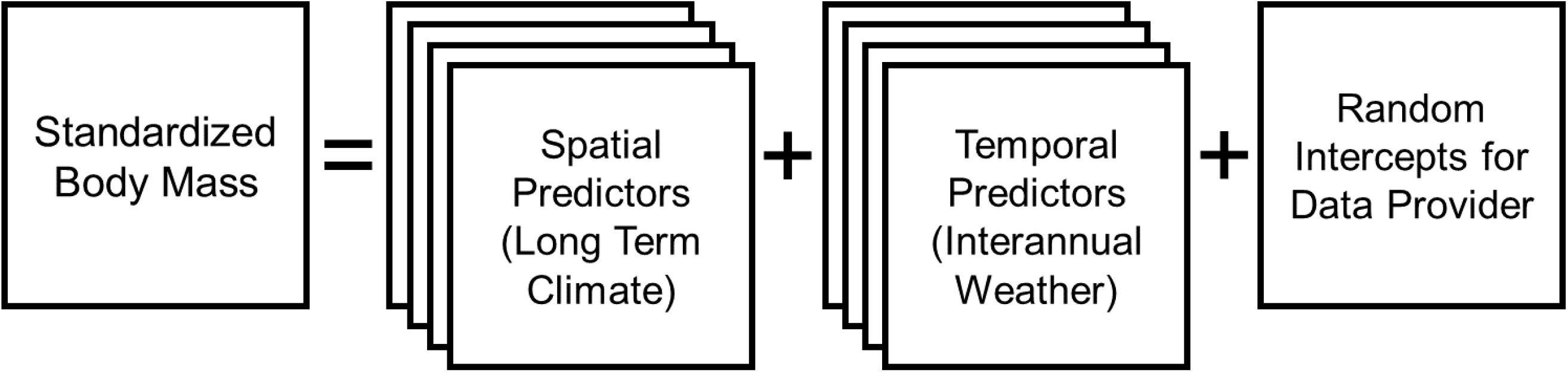
Graphical representation of an individual model. In each model, standardized body mass is a function of the four proxies for the spatial hypotheses, the four proxies for the temporal hypotheses, and random intercepts for each data provider (to account for known differences in protocols for weighing bats). This approach allows tests for the relative contribution of each hypothesis in shaping body size while accounting for the others. In the event that multiple factors interact to influence body size, this approach also allows detection of contributions to body size from multiple hypotheses simultaneously.

**Fig. A2.**
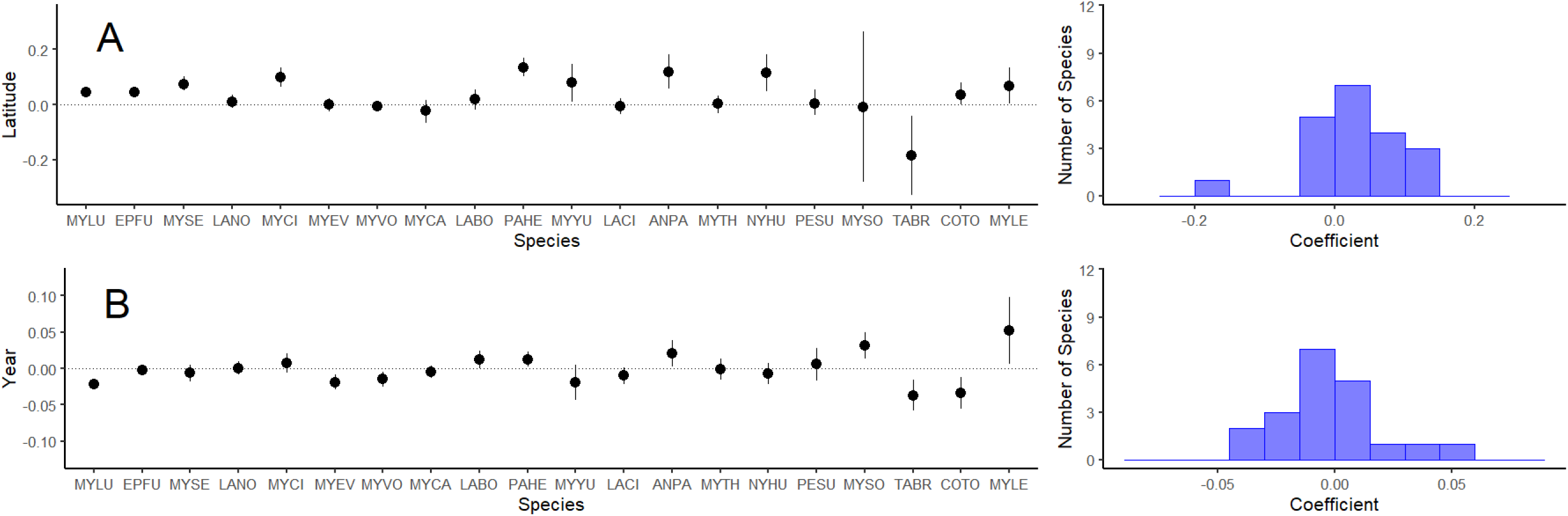
Intraspecific patterns in body mass across latitude and year in 20 species of North American bats. In the left column, we plotted each species’ regression coefficient (points) and 90% credible interval (lines). Points above the dotted line at 0 indicate species with larger masses as the variable of interest (A. Latitude; B. Year) increased. Species are ordered from largest (right) to smallest (left) sample sizes. In the right column, we plotted histograms of the coefficients. Distributions centered on zero indicate no consistent effect of a predictor on body mass, while distributions centered asymmetrically around zero indicate consistent effects. Species tend to be larger at higher latitudes but show no clear pattern of changes in body size over time during our study period.

**Fig. A3.**
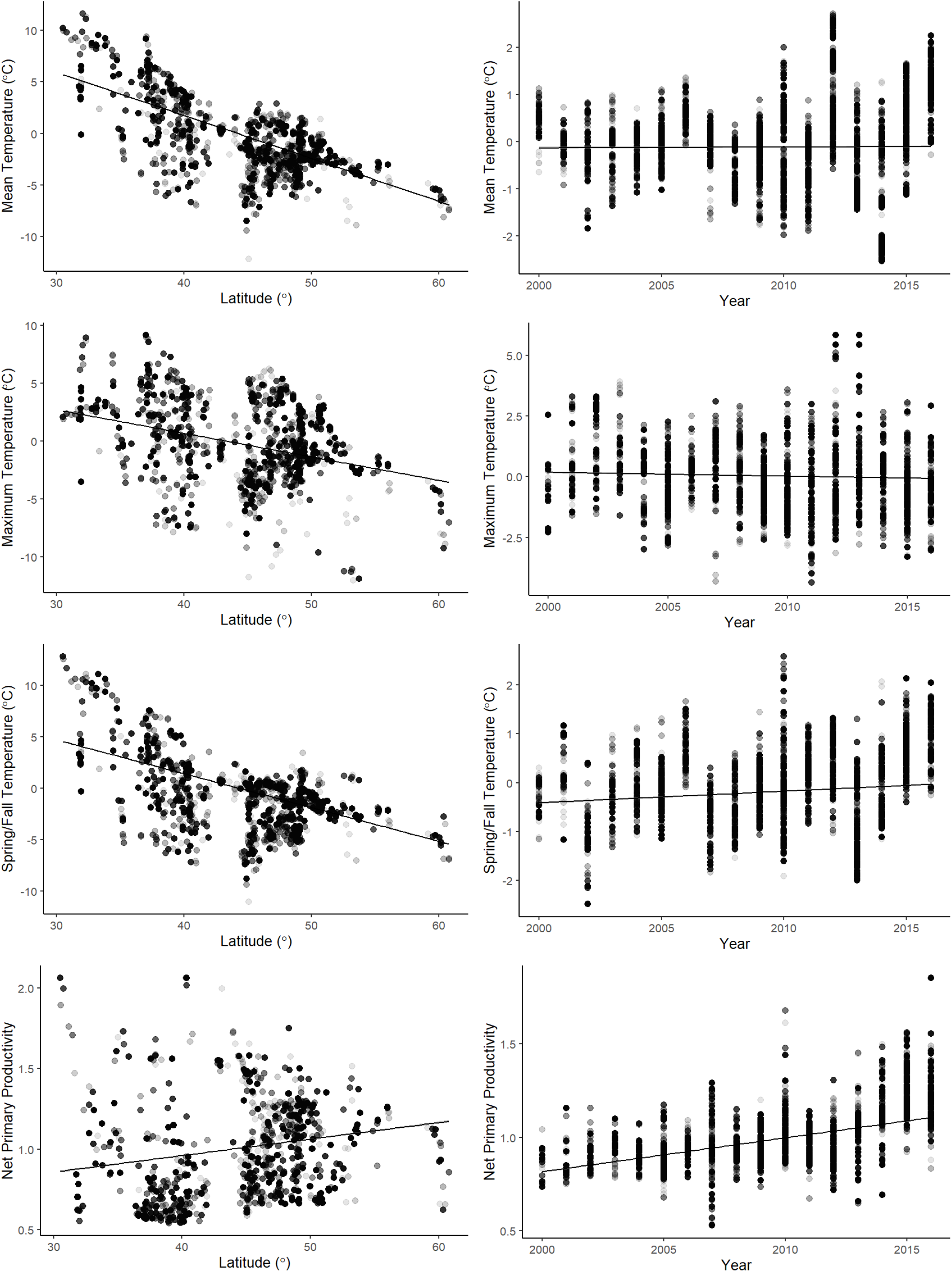
Scatterplots depicting relationships between latitude and variables of interest (left column), and year and variables of interest (right column). Each point represents one capture location, trend lines represent a linear regression of the trend across space or time, and the color of the points represents the number of bats captured at a location (darker points denote more captures). Confidence intervals (95%) are represented by gray ribbons (which are very narrow in all regressions due to large sample sizes). Contrary to expectations, net primary productivity at capture sites increased at higher latitudes (due in part to a large number of bats captured in the arid southwestern United States where primary productivity is low, and the exclusion of winter months when primary productivity is much lower at more northern sites). Following expectations, mean annual temperatures, maximum temperatures, and spring and autumn temperatures decreased as latitude increased. If body size is driven by any of these predictor variables, geographic variation in each of these predictor variables (or some combination thereof) could create a spatial pattern of body mass consistent with Bergmann’s Rule. Across time during our study period, spring and autumn temperatures and net primary productivity increased, while mean annual temperatures and maximum temperatures were relatively constant. Consequently, only spring and autumn temperatures and net primary productivity could lead to an observed trend of shrinking body mass over time during our study. Nevertheless, interannual variation in each of these variables was substantial, so if any of these predictor variables are initiating rapid evolutionary change in body size, our analyses are likely to detect it.

## Notes

### Competing Interest Statement

The authors have declared no competing interest.

